# Single cell transcriptomics reveal temporal dynamics of critical regulators of germ cell fate during mouse sex determination

**DOI:** 10.1101/747279

**Authors:** Chloé Mayère, Yasmine Neirijnck, Pauline Sararols, Chris M Rands, Isabelle Stévant, Françoise Kühne, Anne-Amandine Chassot, Marie-Christine Chaboissier, Emmanouil T. Dermitzakis, Serge Nef

**Affiliations:** Department of Genetic Medicine and Development, University of Geneva, 1211 Geneva, Switzerland; iGE3, Institute of Genetics and Genomics of Geneva, University of Geneva, 1211 Geneva, Switzerland; Université Côte d’Azur, CNRS, Inserm, iBV, France

**Keywords:** Single-cell RNA-Sequencing (scRNA-seq), sex determination, ovary, testis, gonocytes, oocytes, prospermatogonia, meiosis, gene regulatory network, germ cells, development, RNA splicing

## Abstract

Despite the importance of germ cell (GC) differentiation for sexual reproduction, the gene networks underlying their fate remain unclear. Here, we comprehensively characterize the gene expression dynamics during sex determination based on single-cell RNA sequencing of 14,914 XX and XY mouse GCs between embryonic days (E) 9.0 and 16.5. We found that XX and XY GCs diverge transcriptionally as early as E11.5 with upregulation of genes downstream of the Bone morphogenic protein (BMP) and Nodal/Activin pathways in XY and XX GCs, respectively. We also identified a sex-specific upregulation of genes associated with negative regulation of mRNA processing and an increase in intron retention consistent with a reduction in mRNA splicing in XY testicular GCs by E13.5. Using computational gene regulation network inference analysis, we identified sex-specific, sequential waves of putative key regulator genes during GC differentiation and revealed that the meiotic genes are regulated by positive and negative master modules acting in an antagonistic fashion. Finally, we found that rare adrenal GCs enter meiosis similarly to ovarian GCs but display altered expression of master genes controlling the female and male genetic programs, indicating that the somatic environment is important for GC function. Our data is available on a web platform and provides a molecular roadmap of GC sex determination at single-cell resolution, which will serve as a valuable resource for future studies of gonad development, function and disease.

## Introduction

Germ cells are the precursors of the gametes, either sperm or eggs, required for sexual reproduction. GCs are initially bipotential, and their sex-specific fate depends on somatic cues provided by the ovarian and testicular environments, rather than the chromosomal sex they carry (1–3). In mice, primordial germ cells (PGCs) arise in the posterior proximal epiblast around embryonic day (E) 6.25. PGCs rapidly proliferate and colonize the gonads at around E10.5 (4, 5).

In fetal ovaries, GCs enter meiosis asynchronously in a wave from anterior to posterior over about two days, between E12.5 and E14.5 (6–8). This entry into meiosis is considered a hallmark of commitment to oogenesis (9). It is triggered by the expression of the pre-meiotic gene *Stra8* and the meiosis-associated gene *Rec8* and, at the same time, by the downregulation of pluripotency markers such as *Oct4* (*Pou5f1*), *Sox2* and *Nanog* (6, 7, 10). The up-regulation of the *Stra8* and *Meiosin* genes (10, 11) depends on the expression of the transcriptional regulator ZGLP1 that triggers the oogenic fate and stimulates the meiotic program (12). All-trans retinoic acid (ATRA) signaling has been long considered as the inducer of *Stra8* expression and germ cell decision to enter meiosis (13, 14). However, ATRA signaling has recently been excluded as the meiosis-instructing substance in oogonia (15, 16).

In contrast, GCs in fetal testes differentiate into prospermatogonia through a number of distinct, potentially interrelated events that occur asynchronously over a period of several days, but does not involve entry into meiosis (for a review see (17) and (18)). GCs adopting the male fate up-regulate cell-cycle inhibitors such as *Cdkn2b* (19) and are mitotically arrested from E12.5 onward (20). They transiently activate the NODAL/CRIPTO signaling pathway (21–23) and down-regulate pluripotency genes such as *Nanog, Sox2* and *Pou5f1* (24). From E13.5 onward, they begin to express malespecific genes including *Nanos2* (25), *Dnmt3l* (26) and *Piwil4* (27), which ensure normal male gametogenesis by regulating spermatogonial stem cell properties.

Although the cellular origins of oogonia and spermatogonia are well documented (28), numerous questions related to the molecular mechanisms underlying their sex fate and differentiation remain. For instance, the transcriptional programs mediating GC sex determination are incompletely understood, and the essential genes and transcriptional regulators orchestrating such specifications remain poorly defined.

To date, most transcriptional analyses relevant for mouse or human GC sex determination have been conducted using traditional methods such as microarrays or bulk RNA-seq on either whole gonads or isolated GC populations at few selected time points (29–41). These studies, although informative, provided only an average transcriptional summary, masking the inherent variability of individual cells and lineage types and thereby limiting their capacity to reveal the precise dynamics of gene expression during GC sex determination.

To determine the sequence of transcriptional events that are associated with GC commitment and differentiation toward oogenesis and spermatogenesis, we performed time-series single-cell RNA sequencing (scRNA-seq) on developing mouse gonads. We recovered 14,914 GCs from XX and XY gonads across six developmental time points from E9.0 to E16.5, encompassing the entire developmental process of gonadal sex determination and differentiation. We reconstructed the developmental trajectories of male and female GCs, characterized the associated genetic programs, and predicted gene regulatory networks that regulate GC commitment and differentiation.

## Material & Methods

### Transgenic Mice

All work on animals has been carried out in accordance with the ethical guidelines of the Direction Générale de la Santé of the Canton de Genève (experimentation ID GE/57/18). *Tg*(*Nr5a1-GFP*) mouse strain was described previously (42) and has been maintained on a CD1 genetic background.

### Mouse urogenital ridges, testes, ovaries and adrenal glands collection

CD-1 female mice were bred with heterozygous *Tg*(*Nr5a1-GFP*) transgenic male mice. Adult females were time-mated and checked for the presence of vaginal plugs the next morning (E0.5). E9.0 (18±2 somites (ts)), E10.5 (8±2 tail somites), E11.5 (19±4 ts), E12.5, E13.5, E16.5 and E18.5 embryos were collected and the presence of the *Nr5a1*-GFP transgene was assessed under UV light. Sexing of E9.0, E10.5 and E11.5 embryos was performed by PCR with a modified protocol from (43). Posterior trunks (E9.0) and urogenital ridges (E10.5, E11.5) from each sex, XY adrenal glands, testes or ovaries were pooled for tissue dissociation. Posterior trunks, urogenital ridges or adrenal glands were enzymatically dissociated at 37°C using the Papain dissociation system (Worthington #LK003150). Cells were resuspended in DMEM 2%FBS, filtered through a 70 μm cell strainer and stained with the dead cell marker Draq7™ (Beckman Coulter, #B25595). Viable single cells were collected on a BD FACS Aria II by excluding debris (side scatter vs. forward scatter), dead cells (side scatter vs. Draq7 staining), and doublets (height vs. width). Testes and ovaries (from E12.5 to E16.5) were enzymatically dissociated at 37°C during 15 minutes in Trypsin-EDTA 0.05% (Gibco #25300054), resuspended in DMEM 2%FBS and filtered through a 70 μm cell strainer. After counting, 3000 to 7000 single cells were loaded on a 10x Chromium instrument (10x Genomics). Single-cell RNA-Seq libraries were prepared using the Chromium Single Cell 3’ v2 Reagent Kit (10x Genomics) according to manufacturer’s protocol. Each condition (organ, sex and developmental stage) was performed in two biological independent replicates.

### Sequencing

Library quantification was performed using the Qubit fluorometric assay with dsDNA HS Assay Kit (Invitrogen). Library quality assessment was performed using a Bioanalyzer Agilent 2100 with a High Sensitivity DNA chip (Agilent Genomics). Libraries were diluted, pooled and sequenced on an Illumina HiSeq4000 using paired-end 26□+□98 + 8□bp as the sequencing mode. Libraries were sequenced at a targeted depth of 100 000 to 150 000 total reads per cell. Sequencing was performed at the Health 2030 Genome Center, Geneva.

### RNA scope and DAZL immunohistochemistry imaging

For RNAscope experiments, E14.5 XX and XY embryos were dissected in Phosphate Buffered Saline (PBS). Samples were fixed in 4% paraformaldehyde (PFA) overnight at room temperature. Five μm sections were processed for RNA *in situ* detection using the RNAscope 2.0 High Definition-RED Kit according to the manufacturer’s instructions (ACDBio). *Rbm38, Supt6 and Tbrg4* probes were designed by ACDBio. Slides were counterstained with DAPI diluted in the mounting medium at 10 μg/ml (Vectashield, Vector laboratories) to detect nuclei, and with DAZL antibody (cat # GTX89448, Genetex; 1:200) to detect GCs. RNAscope results were examined under a confocal microscope Leica DM5500 TCS SPE on an upright stand (Leica Microsystems, Mannheim, Germany), using an ACS APO 63X oil 1.3 NA objective. The lasers used were diodes laser (405 nm, 488 nm and 532 nm). The microscope was equipped with a galvanometric stage in order to do z-acquisitions.

### Bioinformatic Analysis

#### Data processing with the Cell Ranger package, cell selection and quality controls

Computations were performed at the Vital-IT Center for high-performance computing of the SIB (Swiss Institute of Bioinformatics) (http://www.vital-it.ch). Demultiplexing, alignment, barcode filtering and UMI counting were performed with the Cell Ranger v2.1 pipeline (10x Genomics). Algorithms and versions used are listed in the key resources table. Data were mapped to the mouse reference genome GRCm38.p5 in which the eGFP (NC_011521.1) combined with the bovine GH 3’-splice/polyadenylation signals (Stallings et al., 2002) (NM_180996.1) sequences have been added.

For cell-associated barcode selection, we computed the knee point and the inflection point of the ranked barcodes distribution plot. The local minimum between these points on the density curve (density base R version 3.6.1 function, bw=500, n=4096) of the UMI number per barcode was detected using the quantmod package. This local minimum was used as a threshold for cell-associated barcode selections. When no local minimum could be detected between the two points, the nearest local minimum was used. Quality controls regarding abnormal mitochondrial or ribosomal content, UMI number, detected gene numbers, unmapped reads and putative doublet identification (Scrublet `0.2.1-0`) were performed, but no data filtering was applied as no important anomalies were detected. In total, for gonadal samples, we obtained 92,267 cells. It included 13,060 cells from E9.0, 14,962 cells from E10.5, 16,670 cells from E11.5, 20,285 cells from E12.5, 25,794 cells from E13.5 and 16,994 cells from E16.5. In total, we obtained 61,755 XY cells and 46,010 XX cells. Adrenal glands were collected only in XY embyos and 26,639 cells were collected in total with 7,296, 11,255, 8,822 and 7,609 cells collected at E12.5, E13.5, E16.5 and E18.5 respectively.

#### Gene expression normalization

UMI counts per gene per cell were divided by the total UMI detected in the cell, multiplied by a scale factor of 10,000 and log transformed.

#### Batch correction, clustering, UMAP visualization and germ cells selection

To classify the cell populations present in the developing testis and ovary, we selected all genes detected in more than 50 cells (21,103 genes), log normalized their expression and ran Independent Component Analysis (ICA, Seurat package v2.3.0) on these (100 components). ICA extracts non-Gaussian components from data, allowing a better discrimination of small cell populations in heterogeneous dataset (44). We set the number of ICs to 100 as we expected to have less than 100 different populations in our total dataset. Another advantage of ICA is its good performance with nearly no gene filtering, such as highly variable genes selection.

A neighbor graph corrected for batch between replicates was computed using BBKNN (1.3.5) (45) with default values and n_neighbors_within_batch = 3 and the ICA as input (all components).

Clustering was performed using the Scanpy Leiden method with resolution 1 and UMAP were generated using the Scanpy UMAP method with default parameters. We selected clusters with a strong expression of 10 well-known GCs markers (**Figure S1**).

#### Pseudotime ordering of the cells

To order the cells along a pseudotime, we took advantage of the discrete prior knowledge we have about the embryonic day at which each cell was harvested and generated an ordinal regression model (adapted from (46)) to obtain a continuous pseudotime score reflecting the differentiation status of the cells (Bmrm package v4.1). We trained the model using a tenfold cross-validation schema on the whole expression matrix with a correction for unequal numbers of XX and XY cells. The trained model outputs a weight for each gene according to its ability to predict the embryonic time of each cell in an ordered manner. We then predicted a continuous score for all cells using the weights of the 100 top weighted genes to avoid over fitting of the model.

To determine which genes were important for sex specific progression of the cells along time, we trained two distinct models on XX and XY cells and selected the genes with a significantly higher absolute weight for classification (compared to the complete set of genes). We merged the two lists of genes and obtained 688 unique genes, of which 128 were common to both XX and XY lineages, 266 were specific to XX germ cells, and 294 were specific to XY germ cells (**Fig. S2A** and **B**)

#### Gene regulatory network generation

GRN was generated using pyScenic package (v0.9.14) (47). First adjacencies were generated using the grnboost function (48) with the whole expression matrix of all the germ cells as input and the transcription factor list provided with the package. Next, modules were generated, associating groups of genes to a putative regulator using modules_from_adjacencies function. Finally, the obtained modules were pruned on the basis of the presence or absence of the master regulator binding site footprint in the neighborhood of the TSS of each regulated gene. AUCell was then ran on the obtained regulons to infer its activity within each cell.

#### Regulons and genes hierarchical clustering

Regulons or genes were ordered using ward.d hierarchical clustering on the AUC or expression matrix with Spearman correlation distance. Modules were determined using cutree with k=10 for negative regulons and k=30 for positive regulons (packages dendextend v1.13.4, factoextra v1.0.7, tidyr v1.1.0).

#### Heatmaps and expression curves

Heatmaps were generated using R (packages pheatmap v1.0.12 and heatmaps v1.8.0), smoothing was performed using smoothheatmap function on log-normalized expression levels for genes or AUC enrichment values for regulons. Expression curves were generated using ggplot2 (v3.2.0) geom_smooth function with method “gam” on log normalized expression levels.

#### Spliced and unspliced counts analysis

To generate spliced and unspliced counts data, the velocyto.py script from velocyto (v0.17.8) package was called on each bam file with aforementioned reference genome annotation. Spliced and unspliced counts were normalized using scvelo package (v0.1.19) by dividing all UMI counts of a cell by the total counts in the cell, multiplying the result by the median count number in all cells and applying log1p function. Scatter plots of each specific gene with spliced and unspliced counts were generated using Ms and Mu layers (kNN pooled expression) respectively.

#### Ectopic adrenal germ cells analysis

The analysis was performed using aforementioned steps: log normalization, ICA, neighbor graph, and clustering with the same parameters. GC clusters were selected with the same 10 GC marker genes (see **Supplementary information** for details). For pseudotime ordering of the cells, a model was trained on gonadal cells only and the pseudotime for adrenal GCs was predicted from it.

#### Data and source code availability

GCs single-cell RNA-seq data is available on GEO (accession number GSE136220). Both adrenal and gonadal gene expression data are included in ReproGenomics Viewer (49, 50).

#### Web interface to browse single cell data

To allow biologists and other researchers to explore the GC data we created an interactive website: http://neflabdata.info/. This allows anyone to easily display the expression of any mouse gene in our matrix on a UMAP representation of the GC data. Other metadata (such as sex, time point, or cluster) can also be shown on the UMAP.

Our website uses the Cellxgene data explorer application, which has further functionalities, such as allowing the comparison of the expression profiles of multiple genes, as described in their documentation: https://chanzuckerberg.github.io/cellxgene/posts/gallery. We packaged Cellxgene in a Docker hosted on Google Cloud Platform (GCP) with our GC data H5AD output file from Scanpy. On GCP, we run a virtual machine instance of 2-standard machine type with 2 CPUs and 4-8 GB of memory.

## Results

### Generation of a single-cell transcriptional atlas of germ cell sex determination and differentiation

To generate a gene expression atlas of GCs over sex determination, we used droplet-based 3’ end scRNA-seq (10x Genomics Chromium) of XX and XY gonads from mouse embryos at six developmental stages (E9.0, E10.5, E11.5, E12.5, E13.5, and E16.5) (**Fig. 1A** and **B**). The selected time points cover the entire process of gonadal sex determination and span the emergence and differentiation of the major testicular and ovarian lineages including the gonocytes. For each of the 12 conditions, we generated two independent replicates from different pregnancies and sequenced an average of 10,000 cells. The transcriptomes of the individual cells were sequenced at the depth of ~100,000-150,000 reads/cell. Using ten well-established GC markers, namely *Ddx4, Dazl, Mael, Dppa3, Sycp3*, *Pecam*, *Pou5f1*, *Stra8*, *Dmrt1*, and *Dnmt1*, we identified 14,914 GCs among a total of 107,765 cells (**Fig. S1A-C**) (see also **Supplementary information**). Among GCs, 8,329 were XX (55.9%) and 6,585 were XY (44.1%). This included 56 cells from E9.0, 71 cells from E10.5, 946 cells from E11.5, 4,395 cells from E12.5, 6,633 cells from E13.5 and 2,813 cells from E16.5. The median number of detected expressed genes was higher in GCs than in somatic cells with 5,536 and 4,502 respectively (**Fig. S1D**), consistent with data indicating that GCs undergo pervasive and diverse transcription (51). The generation of this single-cell transcriptional atlas allowed us to further analyze the dynamic, the sex-specific features and the response to the somatic environment during germ cell development.

**Figure 1.**
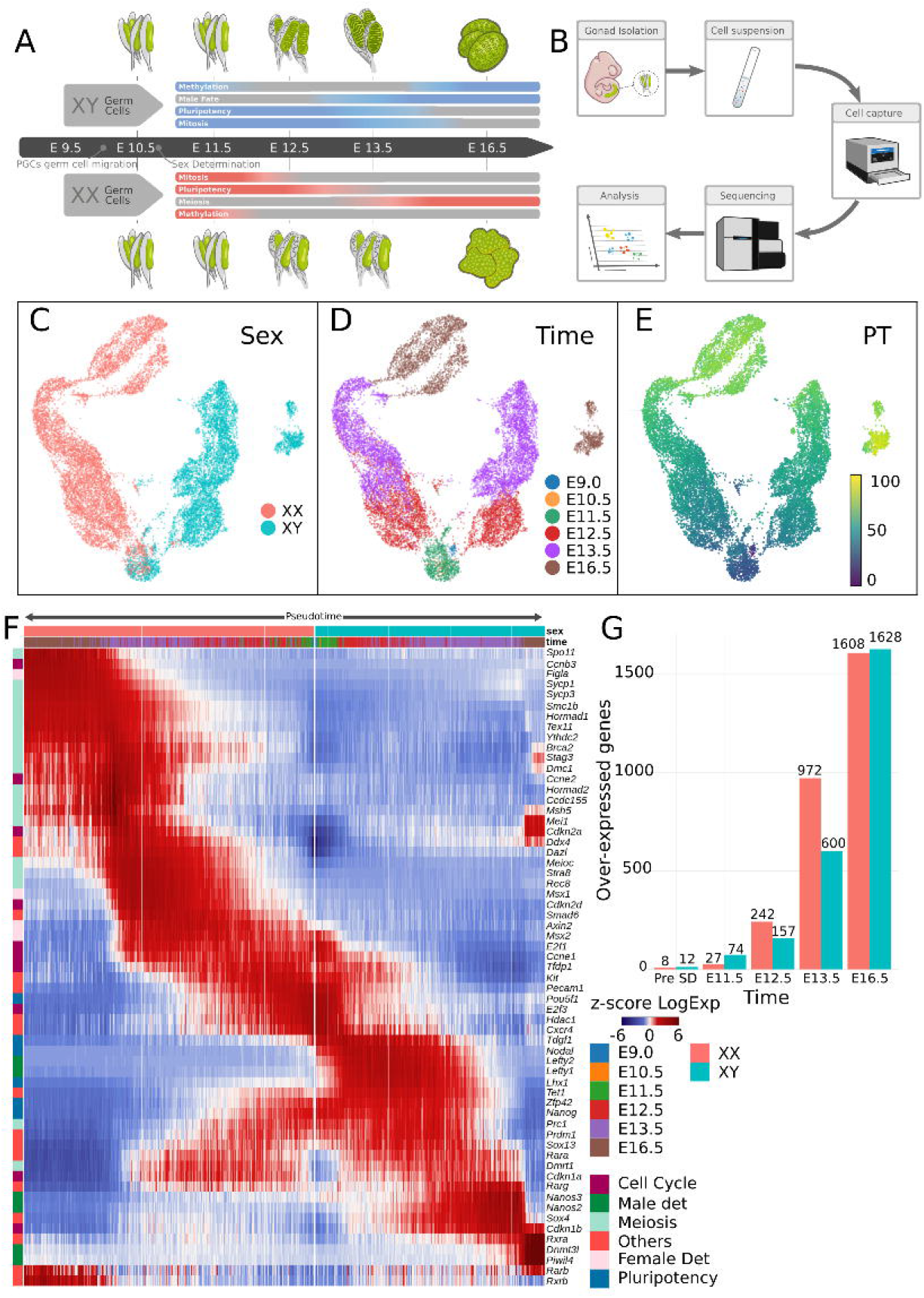
Generation of the germ cell sex determination atlas. (A) Schematic representation of developing testis and ovary highlighting the major events of male and female GC differentiation as well as the time points used in the study. (B) Illustration of the experimental workflow using the 10x Genomics Chromium platform. UMAP projection of 14,914 GCs colored by time (C), sex (D) and pseudotime (PT) going from 0 (E9.5 cells) to 100 (E16.5 cells) (E). (F) Mirror heatmap of 63 known genes involved in GC differentiation. Cells were ordered along a pseudotime. Cells with lowest PT (E10.5) are in the center of the figure and those with highest PT (E16.5) are on the left side for XX cells and on the right side for XY cells. Gene expression was smoothed to reduce dropout effect and obtain a better visualization of expression tendencies (expression scale: log normalized counts normalized per gene). The relevant processes regulated by these genes (cell cycle, male sex determination, meiosis, pluripotency and others) are indicated on the left side of the heatmap. (G) Barplot representing the number of significantly over-expressed genes in XX or XY cells at each stage. The numbers above the bars indicate the number of genes.

### Pseudotime ordering of the cells and differential expression analyses reveals the dynamics of gene expression across XX and XY germ cell sex determination

UMAP projection of the 14,914 GCs permits visualization of the transcriptional landscape (**Fig. 1C-E** and **Material & Methods**). To allow researchers to explore GC expression data, we have created an interactive and user-friendly web site: **http://neflabdata.info/** to easily display the expression of any gene of our dataset on a UMAP representation. Our analysis revealed that, at early stages (E9.0, E10.5 and E11.5), the transcriptomes of XX and XY cells globally overlapped, while more differentiated cells from later stages formed two sex-specific branches (**Fig. 1C** and **D**). We then analyzed how the transcriptomes of XX and XY cells progress during the process of sex determination by ordering the cells along a pseudotime that captures the transcriptional state of each cell. To construct a pseudotime, we used ordinal regression modeling (46, 52) with prior knowledge about the developmental stage of each cell capture (see **Fig. 1E**, **Fig. S1E**, **Material & Methods** and **Supplementary information**). We then represented the smoothed expression of 688 significant genes for two independent sex specific ordinal regression models (**Table S1**) with a double heatmap (mirror heatmap) in which the center represents the earliest cells (pseudotime 0, E9.5 cells) and the extremities represent the lineage endpoints (pseudotime 100, E16.5) of XX GCs (left) and XY GCs (right). The mirror heatmap revealed that XX and XY GCs diverged as early as E11.5, exhibiting dynamic and sex-specific differentiation programs mediated by hundreds of genes (**Fig. S2**). In addition, using 63 well-known genes involved in mouse GC pluripotency, sexual development and differentiation (18, 53), we confirmed that our single-cell data accurately recapitulated known male and female GC specification patterns (**Fig. 1F**).

To assess when sex-specific genetic programs are initiated in XX and XY GCs, we investigated which genes where differentially expressed between these two populations of GCs at each stage of development, retaining only genes with a mean log fold change (LogFC) > 0.25 and an FDR-adjusted p-value < 0.05. Since we had a relatively small number of GCs at E9.0 and E10.5 (56 and 71, respectively), we combined them into a single group called pre-sex determination (pre-SD). As expected, few genes (20) were differentially expressed prior to sex determination (**Fig. 1G**). We then observed a sharp increase in differentially expressed genes (DEGs) with 101 genes at E11.5, whose functional annotations are linked to chaperonine complex and proteasome/proteolysis (see **Tables S2** and **S3** for more detail about DEGs and GO biological process enrichment analysis). Then, 399, 1572 and 3,236 DEGs were identified at E12.5, E13.5 and E16.5, respectively with the XX DEGs associated with meiotic cell cycle, female gamete generation and synaptonemal complex assembly, while the XY DEGs were enriched for RNA splicing and posttranscriptional regulation functions. In XX GCs, a substantial proportion of the DEGs were located on the X chromosome, in particular at E11.5 (**Fig. S1F**). These results are consistent with the gradual reactivation of X-linked genes after E10.5 in XX GCs (54).

Interestingly, E12.5 XY DEGs were also associated to terms such as “axis specification”, “anterior/posterior axis specification”, and “somite rostral/caudal axis specification”. Detected genes associated to this category are *Pitx2, Lhx1, Lefty1, Otx2, Wnt3, Ski, Tdrd5, Tmed2* and include multiple Nodal/Smad2/Smad3 targets (55). Among these genes *Lefty1, Lefty2* and *Pycr2* were upregulated from E11.5 to 13.5 in XY GCs.

Conversely, the BMP pathway target gene *Id1* was found differentially expressed and upregulated in XX compared to XY cells from E11.5 to E16.5. This is consistent with the described activity of Nodal and BMP pathways in XY and XX GCs respectively (21, 56). Taken together, these results indicate that XX and XY GCs diverge transcriptionally as early as E11.5. At this stage, a small set of genes is differentially expressed between XX and XY GCs, but some of them-mostly involved in BMP and Nodal/Activin pathways-mark the first steps of transcriptional differences that will increase throughout cell differentiation (discussed later). Afterwards, the number of differentially expressed genes increases drastically and involves meiosis in XX GCs and multiple biological processes in XY GCs, including RNA processing and pluripotency maintenance.

### Reconstructing Gene Regulatory Networks mediating germ cell sex determination reveals the sequential and modular architecture of germ cell development

The transcriptional state of a cell emerges from an underlying gene regulatory network (GRN) in which the activities of a small number of key transcription factors and co-factors regulate each other and their downstream target genes (47). To comprehensively infer the GRNs at play during XX and XY GC sex determination, we applied the pySCENIC pipeline (47) to our single-cell data. In brief, SCENIC links cis-regulatory sequence information together with scRNA-seq data. It first catalogs coexpression (or repression) modules between transcription factors and candidate target genes and then identifies modules for which the regulator’s binding motif is significantly enriched across target genes; the identified modules are called regulons. Finally, the activity of each regulon is attributed to each cell, using the AUCell method (47). For GCs, we identified 950 regulons (487 positive and 463 negative regulons) containing 11,336 genes (**Table S2**). To compare how XX and XY GCs acquire their sex-specific identity, we selected the 406 positive regulons with binarized AUCell activity in more than 100 cells and containing more than 5 genes, and classified them according to their expression pattern into 30 modules (M1 to M30) along pseudotime (**Fig. 2**). We represented the smoothed regulon expression level of XX and XY GCs with a mirror heatmap. Each module was also functionally annotated via enrichment analyses using GO biological process and Reactome Pathway terms. The most relevant enriched categories are displayed on **Figure 2.** Strikingly, the expression patterns revealed numerous stage and sex-specific regulon profiles, mostly at late developmental stages (late E13.5 and E16.5). Initially, three modules common to both XX and XY gonocytes (M1, M4 and M6, which were cell cycle and splicing associated) were observed at early developmental stages, M1 and M6 persist longer in XY and XX gonocytes respectively. These common modules were superseded sequentially by a handful of transient and overlapping sex-specific regulon modules, M5 (associated to response to BMP, cell cycle arrest and FGF response), M26 (Wnt signaling associated), M27, M29 (associated to meiosis I) in XX GCs, and M2 (RNA splicing and cell cycle associated), M10 (cell junction, cell cycle arrest), M16 and M17 (mRNA processing) in XY GCs. By late E13.5 and E16.5, we noted numerous oogonia-specific like M21, M22 (Calmodulin induced events), and M30 (associated to corticosteroid signaling pathway, Chromatin modifying enzymes and SUMOylation) and spermatogonia-specific modules such as M11 (response to FGF, signaling by RTK), M15, as well as late common modules like M8 (associated to meiosis I, male gamete generation and cell cycle arrest), M9, M13, M14, and M19 (associated with DNA methylation).

**Figure 2.**
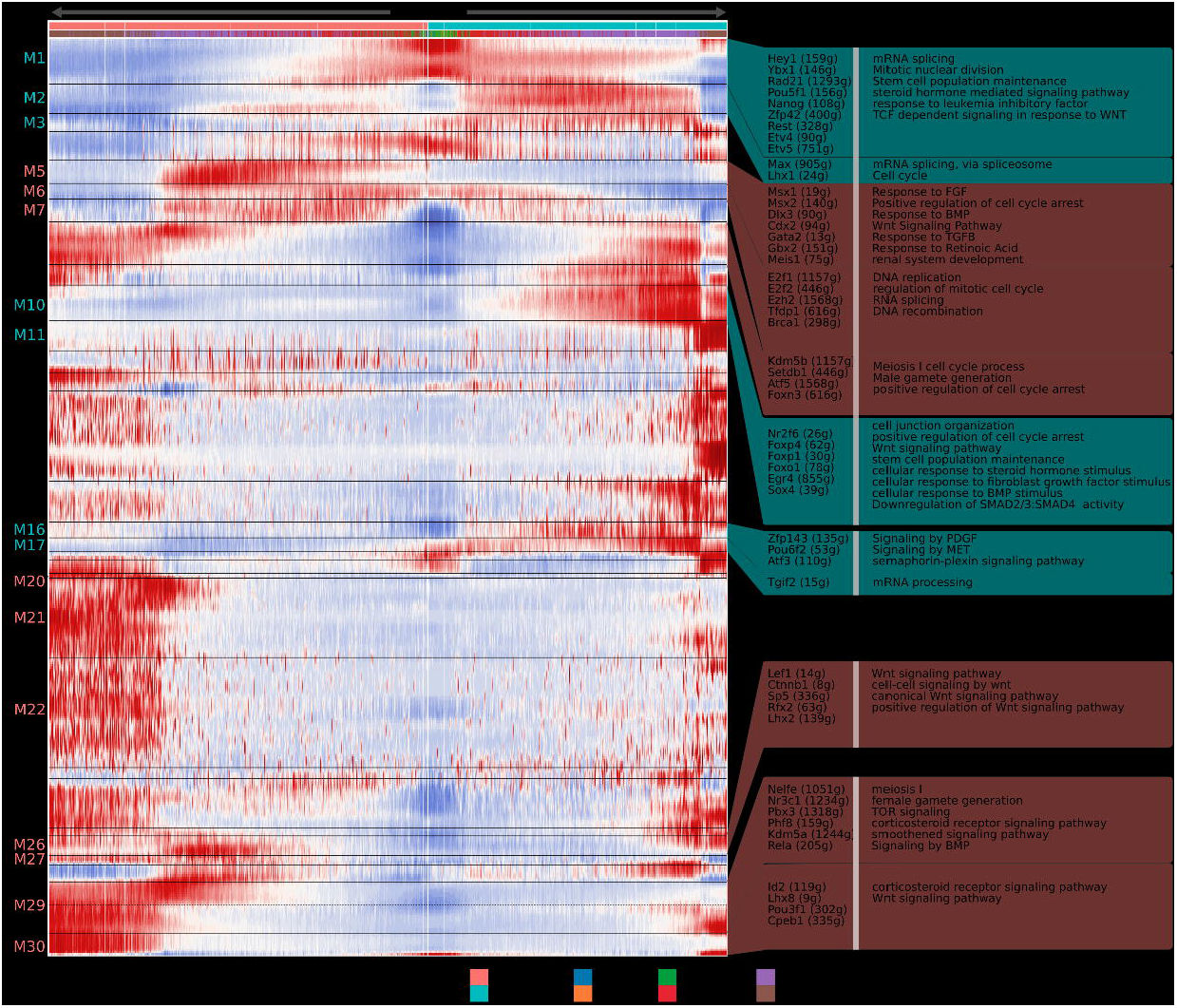
Gene regulation network analysis reveals transient patterns of transcription activation during germ cell sex determination. Mirror heatmap of the 406 regulons with positive association with their master regulator. Regulons were clustered in 30 modules (M1-M30) using hierarchical clustering with Spearman correlation distance. Cells were ordered according to their pseudotime score with lowest score (E9.5) in the center of the figure and highest score (E16.5) on the left side for XX cells and on the right side for XY cells. Boxes on the right display examples of master regulators of interest that are colored by dominant activity in XX (pink) or XY (blue) germ cells. In parentheses is the number of target genes for each master regulator.

We also selected 248 negative regulons with the same criteria and classified them according to their expression pattern into 10 modules (M1 to M10) (**Figure S3**). Negative regulons (-) all displayed activity in opposition to their repressing master regulator and most of them showed a pattern with high activity at early times points, and a progressive downregulation/repression over time (M5 to M10 containing 222 regulons). Among them, regulons of M4(-), M5(-), M9(-) and M10(-) could be distinguished by the onset of their sex-specific repression: regulons from M4(-), M5(-) and M9(-) showed a repression starting earlier (E12.5) in XX cells than in XY cells. We also detected 4 modules formed by 26 regulons with sequential transient expression: M1(-) (associated with meiosis, DNA methylation and response to BMP), M2(-) (positive regulation of cell cycle arrest), M3(-) (DNA methylation) and M10(-) (DNA replication). Among these, target genes belonging to regulons of modules 1 and 10 were transiently repressed specifically in XY GCs, in accordance with the male-specific expression of their 11 master regulators. This implies a possible active repression of the oogonial program in XY GCs by a few master regulators.

Overall, we identified and functionally annotated 654 regulons whose activities were grouped under 40 modules based on their sex-specific and temporal expression. The transient, sequential, and often sex-specific profiles likely represent a sequential/hierarchical organization of regulatory modules required for oogonia and spermatogonia differentiation. The regulons provide a good summary and new insights on the regulation of the events occurring during GC differentiation, for instance, the negative regulation of oogonial fate and the strong presence of mRNA processing associated transcripts in XY GCs.

### RNA splicing is differentially regulated in XY compared to XX germ cells

As mentioned above, we found a strong enrichment of RNA splicing and mRNA processing terms in XY over-expressed E13.5 DEGs. Messenger RNA splicing represents a powerful mechanism to modulate gene expression and is known to contribute to the fine-tuning of cell differentiation programs (57). 51 DEGs are associated with mRNA processing term and their XY-specific or XY-enriched expression profiles between E12.5 and E16.5 are illustrated in **Figure 3A**. The specific expression of genes associated with mRNA processing in XY GCs was confirmed by *in situ* hybridization at E14.5 for the *Tbrg4, Rbm38* and *Supt6* genes (**Fig. 3B** and **C**). *Supt6* codes for a transcription elongation factor which also has a role in mRNAs splicing (58). *Tbrg4* gene encodes a known mitochondrial RNA binding protein that may also have a role in cell cycle regulation (59), while *Rbm38* is coding for an RNA binding protein involved in cell cycle arrest and mRNA splicing (60). RNA-Scope^®^ *in situ* hybridization analysis of these three genes, followed by immunostaining for the DAZL GC marker, reveals that these transcripts were colocalized with the DAZL protein and were specifically enriched in XY GCs, consistent with our scRNA-seq data (**Figure 3C**).

**Figure 3.**
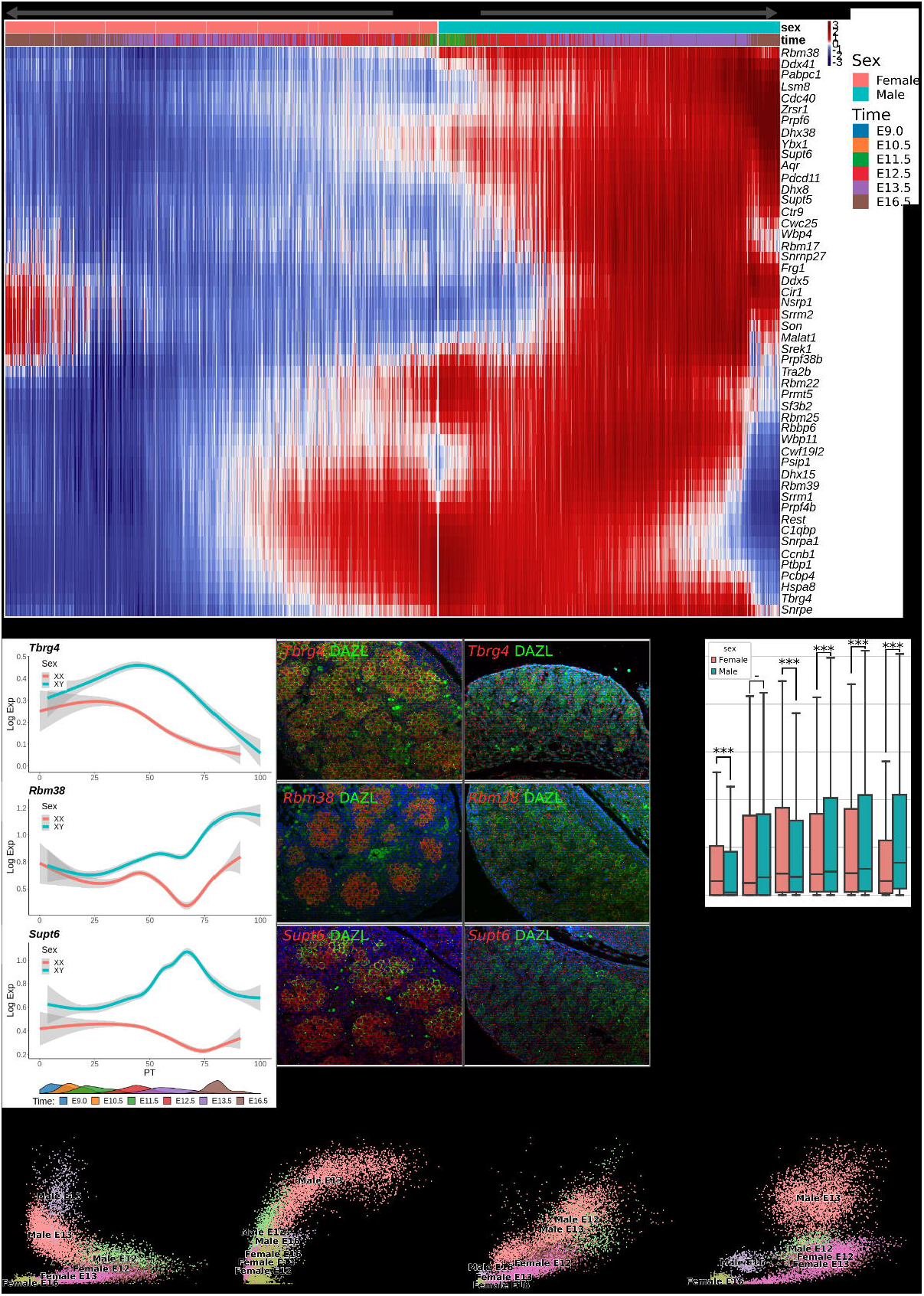
Relative abundance of immature (unspliced) and mature (spliced) transcripts reveals sex- or gene-specific differences in RNA splicing. (A) Mirror heatmap showing the expression profiles of 50 genes associated to the GO term RNA processing found overexpressed in E13.5 XY GCs. Cells and genes were ordered in the same way as in Figure 1F. (B) Expression profiles of *Tbrg4, Rbm38* and *Supt6*, three RNA processing associated genes along pseudotime. Expression levels were smoothed using gam method and 95% confidence interval is figured by grey areas. (C) RNA-Scope analysis followed by immunostaining for the germ cell marker DAZL in E14.5 XX and XY gonads reveals that *Tbrg4, Rbm38* and *Supt6* are enriched in XY GCs. (D) Boxplot showing the mean ratio of unspliced reads over total number of reads for each gene in all cells at a given time. Stars above illustrate the significance level of a Mann-Whitney Wilcoxon test for each comparison: p<0.05:*, p<0.001:**, p<0.0001:***. (E) Scatter plots illustrating a time and sex specific splicing ratio in four selected genes *Ccnd3*, *Bnc2m Tet1* and *Dmrt1*. The X axis represents spliced reads and the Y axis represents unspliced reads. For each plot, a cloud of E13.5 XY GCs show a higher number of unspliced reads compared to all other cells. Expression is in Log normalized counts.

The testis specific expression of RNA processing-associated genes as well as its splicing pattern have already been investigated during adulthood in many species and is known to play a crucial for multiple cellular decision such as mitosis-meiosis transition and stem cell population maintenance (For reviews see (61) and (62)). However, to our knowledge, the presence of a sex specific splicing pattern during embryonic development has not been thoroughly investigated. DEGs overexpressed in XY GCs at E13.5 were associated with terms such as negative regulation of mRNA processing, negative regulation of mRNA splicing and other terms figuring a decreased splicing process. The term “negative regulation of translation” was enriched as well, indicating most likely a global post-transcriptional silencing of gene expression. We therefore formulated the hypothesis that more transcripts should be unspliced in XY GCs at E13.5 compared to XX GCs. Consequently, when looking at each gene in XY and XX GCs, the ratio of intronic/exonic reads should be higher in XY GCs. To test this hypothesis, we counted intronic reads in our dataset and calculated the ratio of intronic counts over total counts (exonic+intronics) for each gene. If the ratio is high, then this indicates that more introns are retained and consequently a reduction in mRNA splicing. We observed a small but highly significant increase in the intronic/exonic ratio in XY GCs (p-value<10^-16^) (**Fig. 3D**) consistent with a reduction in mRNA splicing. This was confirmed at the level of individual genes. Indeed, we have identified genes known for their importance in spermatogenesis, such as *Bnc2, Dmrt1* and *Tet1*, or cell cycle control such as *Ccnd3* whose transcripts have a significantly higher unspliced ratio in XY compared to XX GCs (**Fig. 3E**) (63–66).

Overall, we observed that an important set of RNA processing protein associated transcripts are present in developing XY GCs compared to XX GCs. Moreover, we could infer that RNA splicing is reduced in XY GCs compared to XX GCs, suggesting that this type of post-transcriptional regulation plays an important role in the sex-specific differentiation of the GCs.

### The Nodal/Activin and BMP pathways are important early players in the sexual differentiation of GCs

BMP signaling and the BMP target Zinc Finger GATA Like Protein 1 (ZGLP1) have been reported to be essential for the oogenic fate determination and meiosis entry (12, 56). It has also been shown that the BMP pathway can be repressed at the ligand-receptor level by Nodal/Activin (NA) activity under certain conditions, notably in stem cells (67). In addition, the NA pathway is specifically active in XY GCs from E11.5 to E13.5 and its inhibition allows meiosis of TGC in cultured gonads (21). We therefore investigated the expression of NA and BMP downstream genes in our dataset. As illustrated on **Figure 4A**, we found that NA and BMP target genes are specifically expressed as early as E11.5 in XY GCs and XX GCs respectively and are reciprocally silent in the other sex.

**Figure 4.**
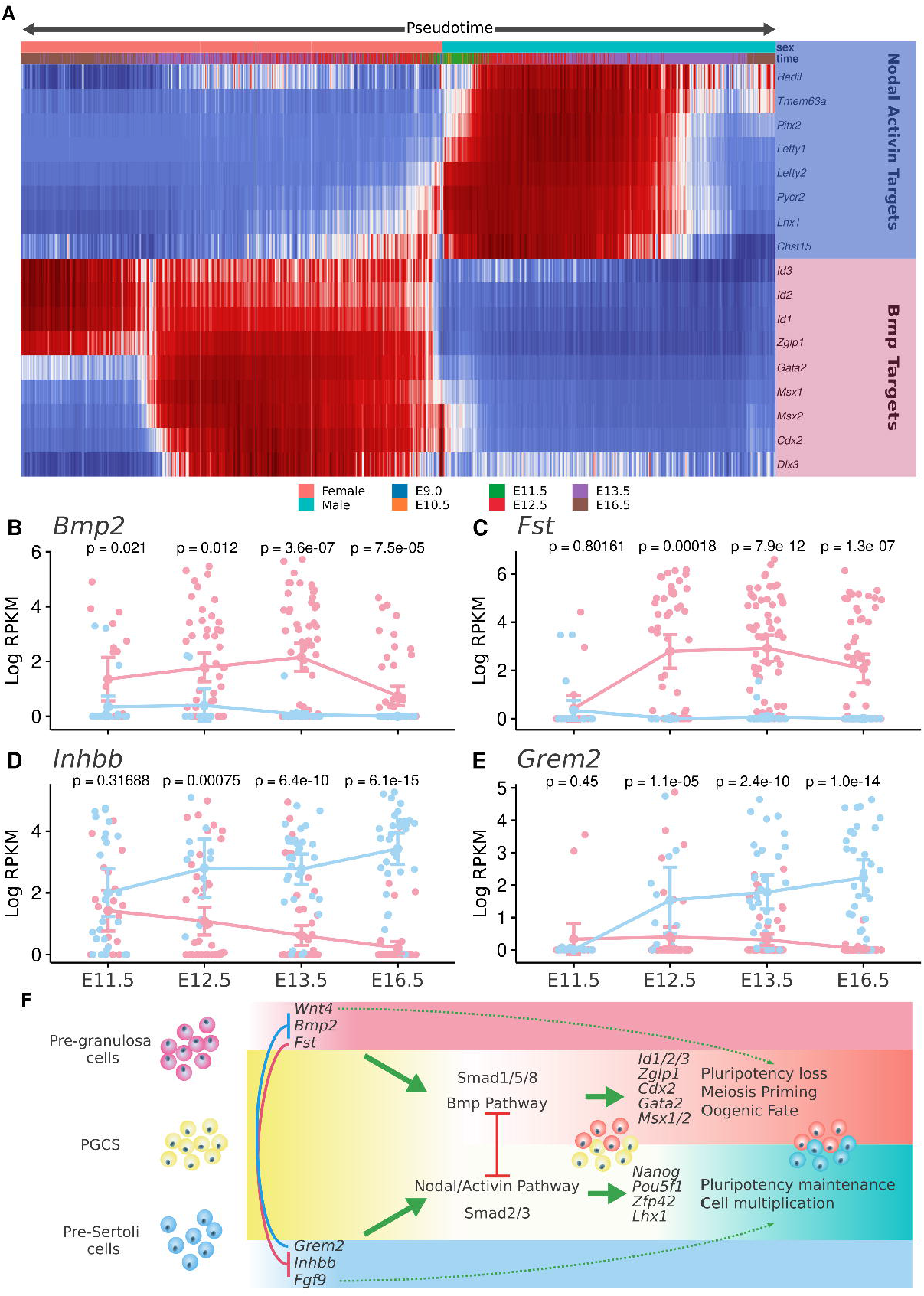
The Nodal/Activin and BMP pathways are important players in the sexual differentiation of GCs. (A) Mirror heatmap representing the expression profiles of ten BMP (pink box) and seven Nodal/Activin (blue box) known target genes. Cells and genes were ordered in the same way as in Figure 1F. Expression profiles of *Bmp2* (B), *Fst* (C), *Inhbb* (D) and *Grem2* (E) in pre-granulosa and (pre-)Sertoli cells at E11.5, E12.5, E13.5 and E16.5. Expression levels are in log RPKM. Above each time point p-values from a Mann-Whitney test comparing the mean of expression of XX and XY cells are displayed. (F) A model of regulation of the BMP and Nodal/Activin pathways in XX and XY PGCs resulting in primary sex differentiation of the GCs as well as meiosis priming for XX ovarian GCs.

In order to identify factors secreted by supporting cells potentially activating or inhibiting the NA and BMP pathways in GCs, we used single-cell RNA sequencing data (scRNA-seq) from somatic XX and XY NR5A1^+^ cells during their differentiation process into granulosa and Sertoli cells (68). We found that *Bmp2* was specifically expressed in pre-granulosa cells as early as E11.5 and not in Sertoli cells (**Fig. 4B**) as described previously (33). We also observed that the gene coding for follistatin (*Fst*), a potent inhibitor of Activin pathway (69), was also overexpressed in pre-granulosa cells from E12.5 onward (**Fig. 4C**). Conversely, we observed a strong sexual dimorphism and overexpression of the beta subunit of activin (*Inhbb*) and the BMP inhibitor *Grem2* in Sertoli cells from E12.5 (**Fig. 4D** and **E**).

Taken together, these findings indicate that GC fate determination in mice occurs as early as E11.5 with the activation of the NA and BMP pathways in XX and XX GCs, respectively. Interestingly, our data suggest that XX and XY supporting cells express and secrete early on factors such as *Bmp2*, *Fst*, *Inhbb* and *Grem2* that either promote one pathway or prevent the activation of the other (**Fig. 4F)**.

**Error! Bookmark not defined.**

### Ectopic adrenal germ cells also enter into meiosis, but numerous major transcriptional regulators of oocyte differentiation are absent or downregulated

While the majority of PGCs migrate toward the gonadal ridges, a small fraction of GCs are lost along the way and end up in extragonadal organs such as the nearby adrenal glands and mesonephroi (70–72). These adrenal GCs (AGCs), irrespective of their genetic sex, have been reported to undergo meiosis, differentiate into oocytes and display morphological characteristics identical to those of young oocytes in primordial follicles before disappearing around 3 weeks of age (71, 72). In the developing adrenal, AGCs are not surrounded by supporting cells such as granulosa cells in the ovary and Sertoli cells in the testis. So, to evaluate how an extragonadal somatic environment affects GC fate, we investigated at the transcriptional level how ectopic adrenal GCs enter into meiosis and commit toward the female fate.

We conducted time-series 3’ single-cell RNA sequencing of developing mouse adrenal cells from E12.5, E13.5, E16.5, and E18.5 XY embryos and identified 312 adrenal GCs based on the expression of the classical GC markers *Dazl, Ddx4* and *Mael* (see **Material & Methods**). Overall, we captured 187 AGCs at E12.5, 92 cells at E13.5, 18 cells at E16.5, and 15 cells at E18.5. The relatively low number of GCs at later stages may reflect the smaller proportion of GCs in the growing adrenal glands or their premature death in this environment. UMAP representation of these 312 XY AGCs combined with our 14,718 gonadal GCs revealed that the transcriptome of XY AGCs partially overlapped with the transcriptome of XX ovarian germ cells (OGCs), suggesting that XY AGCs enter into meiosis and differentiate into oocytes in synchrony with gonadal oocytes (**Fig. 5A**).

**Figure 5.**
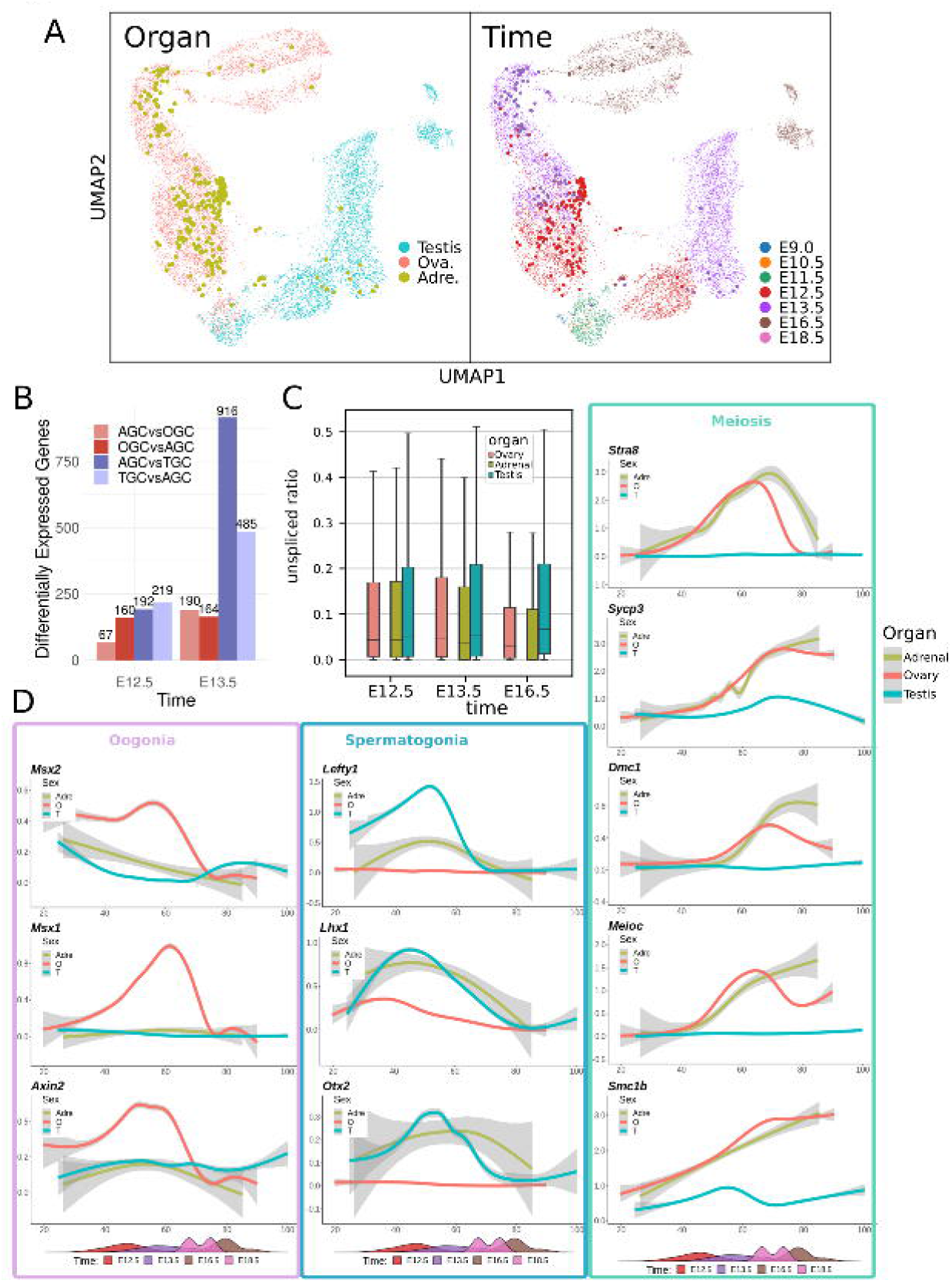
Altered identity and delayed meiosis in adrenal XY germ cells. (A) UMAP projections of 14,718 gonadal cells and 312 ectopic XY GCs developing in the adrenal colored by organ (left) and time (right). Adrenal GCs are represented in green, with larger dots. (B) Number of differentially over-expressed genes for each comparison adrenal GC (AGC) vs ovarian GC (OGC,) AGC vs testicular GC (TGC) and reciprocally at E12.5 and E13.5. (C) Boxplot representing splicing ratios observed in AGC, TGCs and OGCs. (D) Expression profiles of selected genes involved in meiosis, oogonia and spermatogonia differentiation process. The solid line represents the smoothed expression curves of the gene in the cells ordered by pseudotime, and the fade band is the 95% confidence interval of the model.

To investigate further the similarities and differences between AGCs, testicular germ cells (TGCs) and OGCs, we ran differential expression analysis between these three cell populations at meiosis onset (E12.5 and E13.5) (**Table S4**). We found that the comparison between AGCs and TGCs yielded more differentially expressed genes than the comparison between OGCs and AGCs, in particular at E13.5 (**Fig. 5B**). Enrichment analysis of GO terms revealed that terms associated with meiosis are enriched in genes over-expressed in AGCs compared to TGCs while terms associated with splicing and mRNA processing are enriched in TGCs over-expressed genes (**Table S5**). Likewise, splicing ratios were very similar between AGCs and OGCs, confirming that AGCs have overall transcriptional characteristics comparable to OGCs (**Fig. 5C**).

Despite these strong transcriptomic similarities, some significant differences could nevertheless be observed between AGCs and OGCs. In general, meiosis-related genes showed similar patterns and levels of expression between AGCs and OGCs. This is the case for *Stra8, Sycp1, Sycp3, Sync3, Spo11, Ccdc155, Dmc1, Mei1, Mei4, Meioc, Hormad1, Hormad2, Msh5, Tex11, Prdm9, Zglp1* and *Smc1b* (**Fig. 4D** and **Fig. S5**). However, a notable exception was *Rec8*, whose expression is reduced in the adrenals compared to ovarian GCs (**Fig. S4A**). These results confirmed published data that meiosis is not significantly affected in ectopic AGCs (71, 72). However, we found numerous key female genes exhibiting either downregulation or a testis-like profile, for example, genes involved in the WNT-β-catenin pathway (*Axin2, Lef1*, and *Sp5*) and BMP target genes transcription factor genes such as *Gata2, Id1, Id2, Id3, Cdx2, Smad6, Dlx3*, *Msx1* and *Msx2*, the cell cycle gene *E2f1*, and the oocyte-specific basic helix-loop-helix transcription factor gene *Figla* (**Fig. 4D** and **Fig. S4A and B**). Finally, we found that most genes involved in male GC fate were not upregulated in AGCs with few notable exceptions including the retinoic receptor gene *Rara* and the male GC regulator gene *Ctcfl*. Some other genes such as the NODAL target genes *Lefty1, Lefty2, Pitx2, Lhx1* (a spermatogonial stem cell self-renewal gene) and *Otx2* and the male fate marker *Nanos2* showed an intermediate expression between OGC and TGC level. (**Fig. 4D** and **Fig. S4B**). Overall, these results indicated that ectopic adrenal XY GCs enter into meiosis at around the same time as ovarian XX GCs (and unlike their XY counterparts in the testis), but numerous genes related to both the female and male genetic programs were misregulated.

## Discussion

This study represents the first large-scale scRNA-seq analysis of XX and XY GCs throughout the gonadal sex determination and differentiation process. The large number of individual GCs profiled, allowed us to reconstruct a near continuous representation of changes in gene expression occurring during the process of gonadal differentiation, including the mitosis-to-meiosis switch in GCs in developing ovaries, and spermatogonial commitment and differentiation in fetal testes. This represents a major advance beyond previous work and has broad implications for studies in GC development, sex determination and the etiology of human GC diseases. We also provide a user-friendly website to display the expression of any gene expressed in XX and XY GCs during gonadal sex determination using a UMAP representation at **http://neflabdata.info/.**

How do GCs commit to and acquire sex-specific fates during differentiation of the gonad into a testis or an ovary? From our time course scRNA sequencing profiling of six developmental stages encompassing gonadal sex differentiation, we could observe sex specific transcriptional patterns as early as E11.5. At this stage, part of this differential expression can be explained by the X chromosome reactivation occurring in XX GCs. However, we found that the Nodal/Activin and BMP pathways are important early actors in the sex-specific differentiation of GCs with the early and sex-specific expression of multiple target genes of the Nodal/Activin pathway expressed from E11.5 onward in XY GCs as well as numerous target genes of the BMP pathway upregulated in XX GCs. In addition, we found that RNA splicing is differentially regulated with increased unspliced mRNAs in XY GCs around E13.5 potentially promoting male GC fate, regulating mitotic arrest, inhibiting meiotic entry and/or antagonizing the female GC fate.

### Transient, sequential – and often sex-specific – profiles of regulatory modules mediating oogonia and spermatogonia differentiation

Here we uncover the gene regulatory networks (GRNs) driving sexual differentiation of the GCs in a comparative way. We have comprehensively constructed the GRNs for XX and XY GCs during the process of sex differentiation. We found that specific combinations of transcription factors drive sex-specific transcriptomes in GCs and that the gene regulatory circuitry mediating GC sex determination is composed of 654 positive regulons that can be grouped into 40 modules, each of them exhibiting transient, sequential and often sex-specific profiles. The fact that regulons are grouped into modules displaying transient sex-specific profiles suggests a sequential organization of regulatory modules that work together to fine-tune the interrelated cellular events that lead to XX and XY GC differentiation. The master regulator genes present in each specific module may not be controlling a single cellular event, but instead a combination of overlapping sex-specific developmental processes including mitotic arrest, prospermatogonia commitment and *de novo* methylation for XY GCs, as well as entry into meiosis, and suppression of pluripotency genes for XX GCs.

The GRN inference analysis also provides an opportunity to identify new critical master regulators of GC sex determination. While various master regulators have already been implicated in playing a key role in pluripotency and GC sex-specific commitment and differentiation, such as *Dazl, Pou5f1, Dmc1, Rec8, Stra8, Nodal, Nanos2*, and *Dnmt3l*, our analysis predicted more than 800 positive and negative regulons (**Fig. 2, Fig. S3** and **Table S2**), including a large set of new potentially critical regulators of GC commitment and differentiation, for example RAD21, YBX1, EGR4, ETV4, ETV5, KDM5A, KDM5B, NR3C1 and PHF8. These predictions provide an important framework and guide for future experimental investigation.

### Messenger RNA processing and intron retention as post-transcriptional regulation in developing mammalian germ cells

Another level of control of gene expression and function lies in RNA maturation, stabilization and degradation (57). We observed a global post-transcriptional silencing of gene expression in XY GCs around E13.5 potentially involved in repressing premature meiosis entry, regulating mitotic arrest and promoting male GC fate. Not only we found numerous DEGs overexpressed in XY GCs associated with terms such as negative regulation of mRNA processing and splicing but also determined that RNA splicing is globally reduced in XY GCs compared to XX GCs. We found a large variation in the ratios of unspliced/spliced transcripts between XX and XY GCs suggesting that intron retention is another differential feature of XY GCs, allowing them to adopt the appropriate sexual fate through sex-specific differential expression. Finally we provide example of genes known for their importance in spermatogenesis, such as *Bnc2*, *Dmrt1* and *Tet1* (63–65), or cell cycle control such as *Ccnd3* (66) which all displayed increased levels of unspliced mRNAs in XY GCs. Intron retention has been shown to be a prominent feature of the meiotic transcriptome of mouse spermatocytes (73). It can either favor accumulation, storage, and timely usage of specific transcripts during highly organized cell differentiation programs or cause transcript instability at specific developmental transitions (73–77). The temporal or sex-specific variation in intronic retention appears to be surprisingly frequent during the process of GC sex determination and may ensure proper and timely expression of selected transcripts. Full length isoform single cell sequencing with long reads would further illuminate and validate our splicing results and the importance for the male specific differentiation of GCs (78, 79).

### Nodal/Activin and BMP show opposite activity patterns in XX and XY GCs and may be a kick-starter of germ cell sexual differentiation

Historically, STRA8 has been described as the only gatekeeper that engages the meiotic program in developing female GCs (10). The onset of *Stra8* expression in GCs of the developing ovary and its lack of expression in GCs of the developing testis led to a search for the presence of the female “meiosis-initiating substance” (MIS) or male “meiosis-preventing substance” (MPS) (17, 80). While RA has emerged as a potential MIS (13, 14), recent reports revealed that female GCs enter meiosis normally even in the absence of RA signaling (15, 16, 81). Recently Nagaoka et al. showed that BMP signaling and its downstream transcriptional regulator ZGLP1 were essential for the oogenic fate, with RA signaling assisting in the maturation of the oogenic program as well as to the repression of the PGC program (12). On the other hand, it is interesting to note that many BMP target genes such as *Gata2, Id1, Id2, Id3, Msx1, Msx2, Cdx2* were not over-regulated in AGCs and have an expression profile closer to TGCs (**Fig. S5**). This suggests that a strong BMP signalling is dispensable for meiosis initiation and specification of female GC fate, at least in the adrenal gland (**Fig. 5A**). It also suggests that upregulation of *Zglp1* in AGCs requires little or no BMP activity. However, as AGCs do not survive later in adrenal gland, we can’t exclude BMP signaling may be important for cell survival and complete oogenic differentiation.

There are clear evidence that the somatic environment of the embryonic gonad, in particular the supporting lineage, influence GC sex determination through the secretion of factors that act either as promoting or inhibiting factors (17). Investigating the expression profile of developing Sertoli cells and granulosa cells, we identified several sex determining cues secreted by these supporting cells that promote or inhibit the BMP and Nodal/Activin signaling pathways. On one side, the granulosa cells promote the BMP pathway through the expression of *Bmp2* and repress Activin activity via the expression of *Fst*. Conversely, Smad2/3 activity, which is intrinsically triggered by GCs by Nodal expression is reinforced by *Inhbb* which is expressed by Sertoli cells, which also repress the BMP pathway through the expression of *Grem2*. GREM1 and GREM2 are members of the DAN family of BMP inhibitor that bind BMPs and prevent them from associating with their receptors (82, 83). Both *Grem1* and *Grem2* have been shown to be expressed in the postnatal ovary and to have roles in regulating follicular development (84–86). Whether GREM2 has or not an MPS activity in the developing testis requires further investigations.

### Comparing adrenal and gonadal germ cells development provides a tool to investigate the importance of the somatic environment in the process of germ cells differentiation

By comparing the transcriptome of adrenal and gonadal GCs, we have been able to investigate how GCs respond to three different somatic environments: the adrenal, ovarian and testicular environments. It allowed us to also investigate whether the gene regulatory circuitries mediating GC sex determination, composed of 950 regulons, are interconnected or act independently. The dynamic expression pattern of key marker genes of meiosis is strikingly similar in both AGCs and OGCs, suggesting that the initiation and maintenance of meiosis proceeds relatively normally in AGCs, despite their XY karyotype. However, we also observed a significant alteration in the expression of some key female master regulator genes as well as upregulation of some male specific genes, indicating that the somatic environment in the adrenal gland cannot completely support a female fate for these gonocytes. In particular, we observed a lack of upregulation of genes involved in the canonical WNT/β-catenin signaling pathway (*Axin2, Lef1*, and *Sp5*), suggesting that GCs in this environment are unable to respond to WNT signals, or to express their receptors (87–89). The expression of other genes downstream of the BMP or NA signalling pathways such as transcription factors *Id1*, *Id2*, *Id3*, *Dlx3*, *Gata2*, *Msx1* and *Msx2* and *Cdx2* (90) as well as the oocyte-specific basic helix-loop-helix transcription factor gene *Figla* also displayed significant alteration. The findings that downstream genes for both the WNT and BMP signaling pathways are not turned on in AGCs (with the notable exception of *Zglp1*) suggest that these two pathways may be dispensable to initiate meiosis and the female GC fate. Interestingly, the absence of *Msx1* and *Msx2* expression may explain why *Rec8* expression, but not other meiotic genes such as *Stra8*, is blunted in adrenal GCs (**Fig. S4B**). Based on our GRN analysis, we found that both MSX1 and MSX2 are predicted to positively regulate *Rec8* expression.

The large majority of male-specific master regulators are not expressed in XY AGCs with few notable exceptions such as the NA responsive genes *Lefty1*, *Lefty2* and *Lhx1* suggesting partial suppression of the NA activity in XY AGCs. Other genes normally expressed in TGCs such as *Nanos2, Rara* and *Ctcfl* are also expressed in AGCs supporting the idea that overall, the adrenal environment does not provide all the necessary signal(s) required to commit GCs to oogenesis, resulting in AGCs characterized by an altered ovarian identity despite meiosis entry.

Compiling single cell transcriptomes from mouse GCs at six developmental stages during the process of sex determination, both in gonadal and extragonadal tissues, allowed us to provide a comprehensive insight into the sex-specific genetic programs and gene regulatory networks that regulate GC commitment toward a male or female fate. As such, we have created a valuable and rich resource for the broad biology community that should provide insights into how this fundamental decision impacts the etiology of infertility and human gonadal GC tumors, two of the main clinical consequences of defects in GC sex determination.

## Supporting information

Table S1 - Ordinal regression selected genes

Table S2 - Differentially expressed genes XX vs XY

Table S3 - GO enrichments

Table S4 - Differentially epressed genes AGC vs Gonadal GC

Table S5 - Go enrichment AGC DEGs

## Abbreviations

AGC: Adrenal Germ Cell
GC: Germ cell
OGC: Ovarian Germ Cell
TGC: Testicular Germ Cell
scRNA-seq: Single-cell RNA-Sequencing
DEG: Differentially Expressed Gene

## Acknowledgments

This work was supported by grants from the Swiss National Science Foundation (grants 31003A_173070 and 51PHI0-141994) and by the Département de l’Instruction Publique of the State of Geneva (to S.N.). We thank Luciana Romano and Deborah Penet for the sequencing, Christelle Borel (GEDEV department, University of Geneva) for her advice and help with 10X technology, Cécile Gameiro and Gregory Schneiter (Flow Cytometry Facility, University of Geneva), the team of the Animal Facility (Faculty of Medicine, University of Geneva), Julien Prados (Basic Neuroscience, University of Geneva) for his help for pseudotime computation and Valentin Durand Graphic Design for help with artwork. We thank also Andy Greenfield, the members of the Nef and Dermitzakis laboratories for helpful discussion and critical reading of the manuscript.

## Author Contributions

Conceptualization, S.N.; data generation and investigation, C.M., Y.N., P.S., A.A.C., I.S. and F.K.; Formal Analysis and Data Curation, C.M, C.M.R. and P.S.; Writing – Original Draft, S.N., C.M., and M.-C.C.; Funding Acquisition, S.N., and E.T.D.; Resources, S.N., and E.T.D.; Supervision, S.N., and E.T.D.

