## Supplementary Figures and Information for "Single cell transcriptomics reveal temporal dynamics of critical regulators of germ cell fate during mouse sex determination"

### Supplementary information

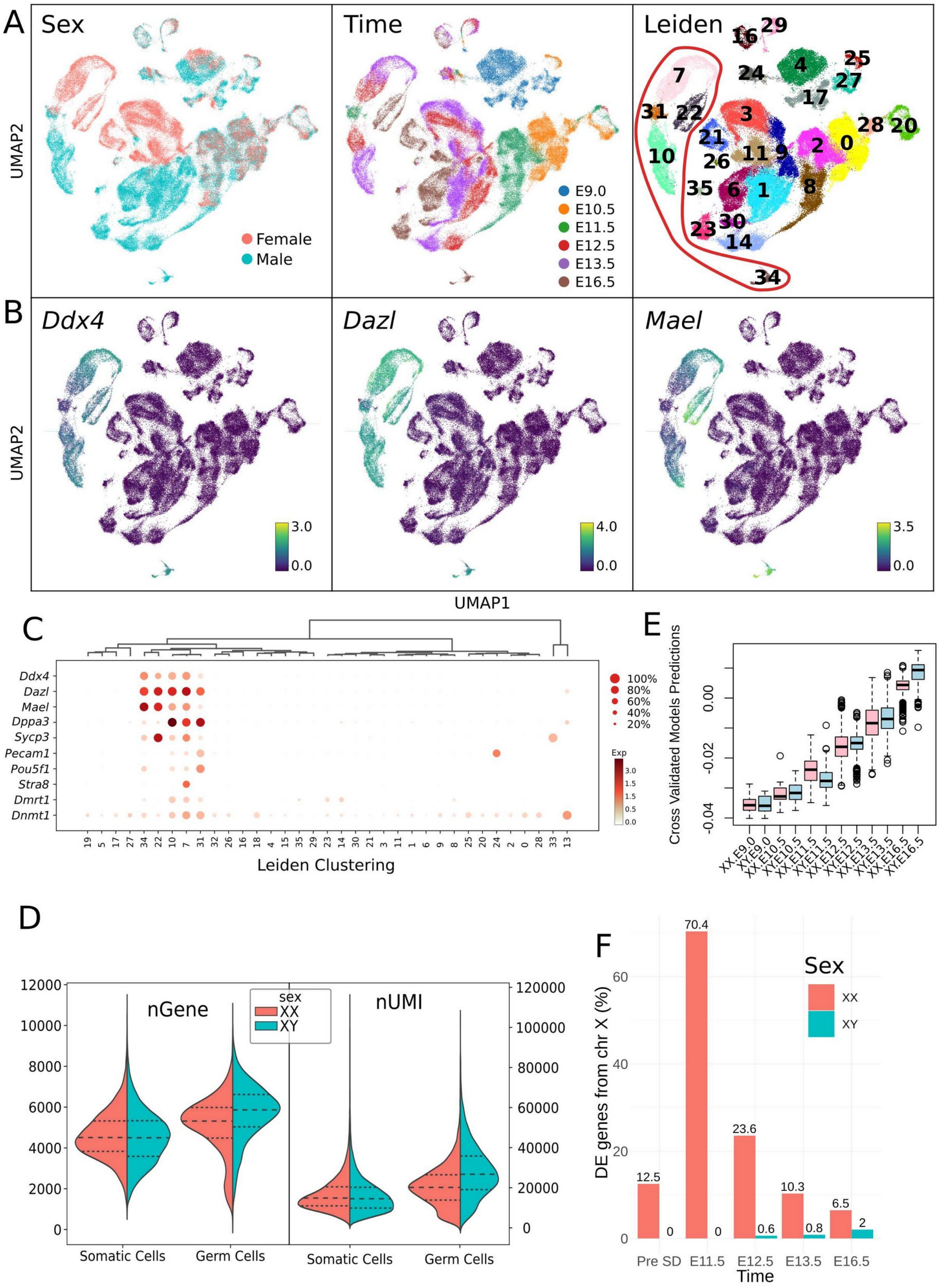


**Figure S1**

**Legend Figure S1** (related to Figure 1)**. Mouse sex determination atlas.** (A) UMAP projection of the 92,267 gonadal cells colored by sex, embryonic days and cluster. The nine germ cell clusters are surrounded by a red line. (B) UMAP projection of the cells colored by expression of the germ cells markers *Ddx4, Dazl* and *Mael*. Expression scale: log normalized counts. (C) Dotplot representation of the expression of germ cells marker genes in each cluster. The size of the node represents the percentage of cells in the cluster expressing the gene and the color indicates the level of expression (log normalized counts normalized per gene). Cells from clusters 34, 22, 10, 7 and 31 are germ cells and will be used for the subsequent analysis. (D) Violin plots of the number of UMIs and detected genes in germ cells against somatic cells, by sex. Dashed lines represent median an dotted lines upper and lower quartiles. (E) Boxplot visualization of the score of the cells in the ordinal regression model for each time and sex. Solid lines represent the median, box limits are the upper and lower quartiles. The pseudotime ordering is obtained from this score scaled from 0 to 100.

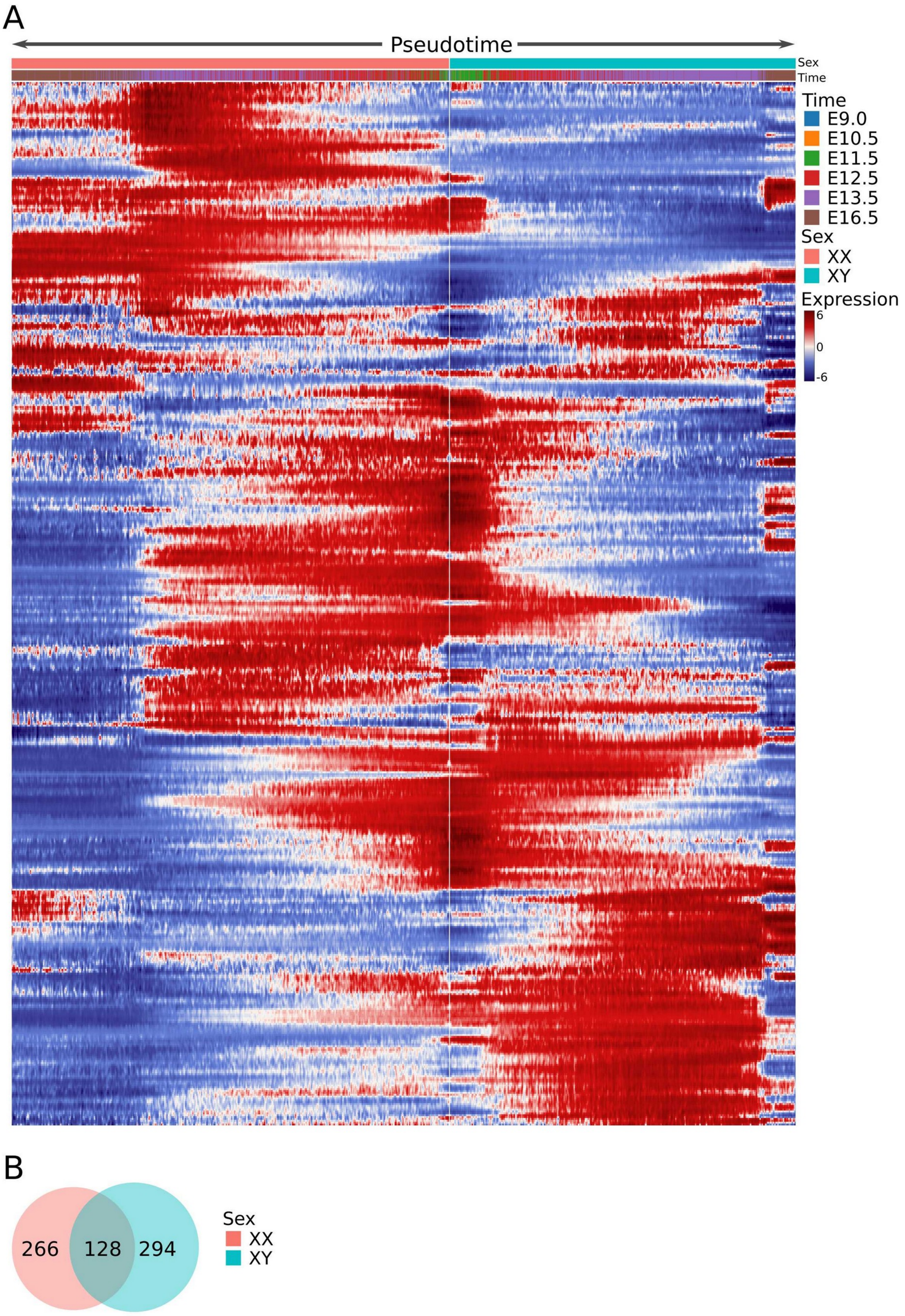
A
Pseudotime
Sex
Time
Time
E9.0
E10.5
E11.5
E12.5
E13.5
E16.5
Sex
XX
XY
Expression
6
0
-6
B
266
128
294
Sex
XX
XY

**Figure S2**

**Legend Figure S2** (related to figure 1)**. Genetic programs mediating germ cell sex determination.** (A) Expression heatmap of 688 genes involved in germ cell transition from E9.0 to E16.5 according to ordinal regression models applied to XX and XY cells separately. Similar gene profiles were hierarchically grouped together using Spearman correlation distance. Cells were ordered using a pseudotime score generated with ordinal regression modeling, expression was smoothed to reduce dropout effect and obtain a better visualization of expression tendencies (expression scale: log normalized counts normalized per gene). Cells with lowest score (E9.0) are in the center of the figure and those highest scores (E16.5) are on the left side for XX cells and on the right side for XY cells.(B) Venn diagram of the number of selected genes.

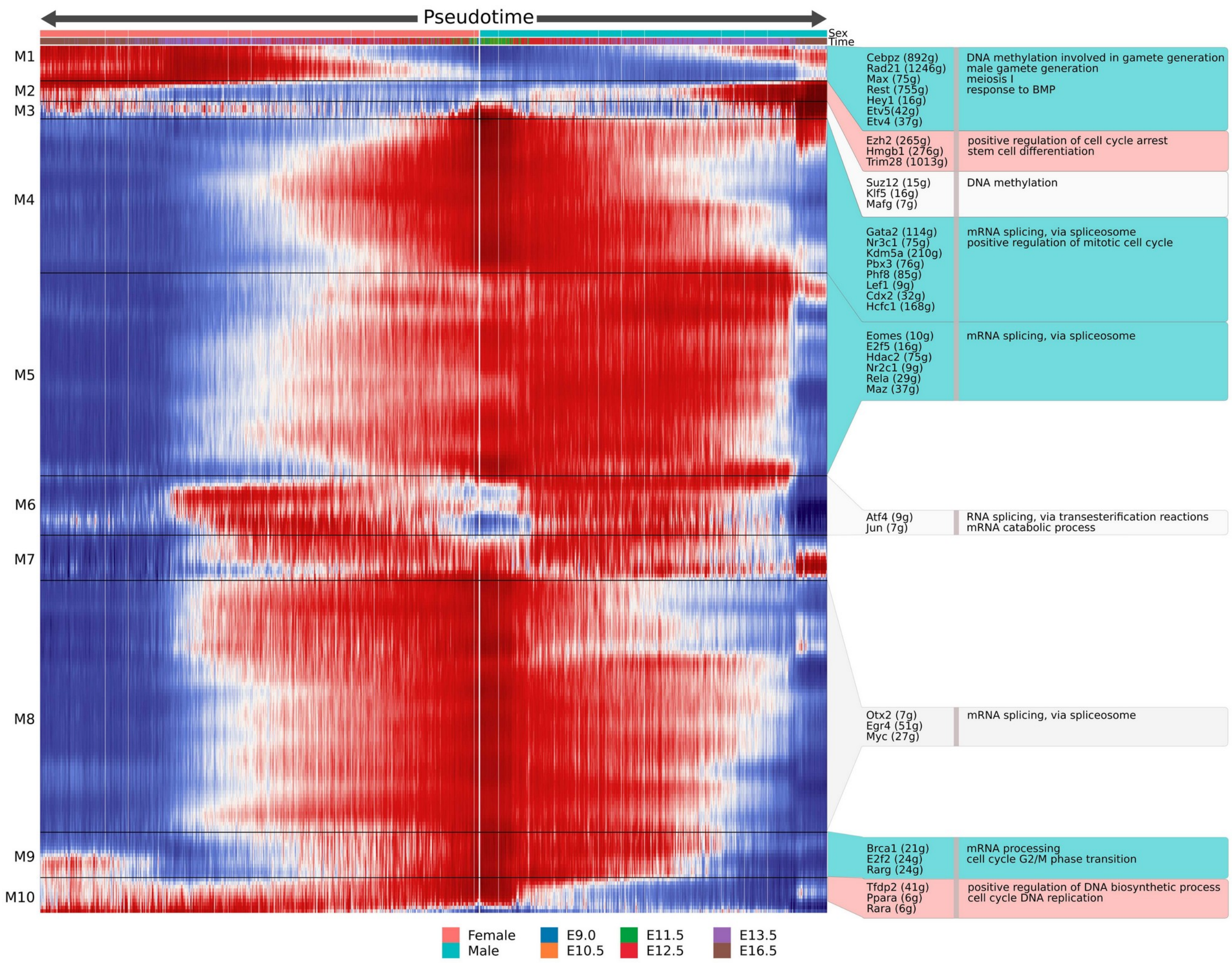


### Figure S3

**Legend Figure S3** (related to figure 2)**. Gene regulation network analysis with negative regulation associations** A. Heatmap of the 303 regulons with a repressive association with their master regulator. Regulons were clustered in 10 modulons using hierarchical clustering with Spearman correlation distance. Cells were ordered using a pseudotime score generated with ordinal regression modeling, expression was smoothed to reduce dropout effect and obtain a better visualization of expression tendencies (expression scale: log normalized counts normalized per gene). Cells with lowest score (E10.5) are in the center of the figure and those highest scores (E16.5) are on the left side for XX cells and on the right side for XY cells. Boxes on the right display examples of master regulators of interest that are colored by dominant activity in XX (pink) or XY (blue) cells. In brackets is the number of target genes for each master regulator.

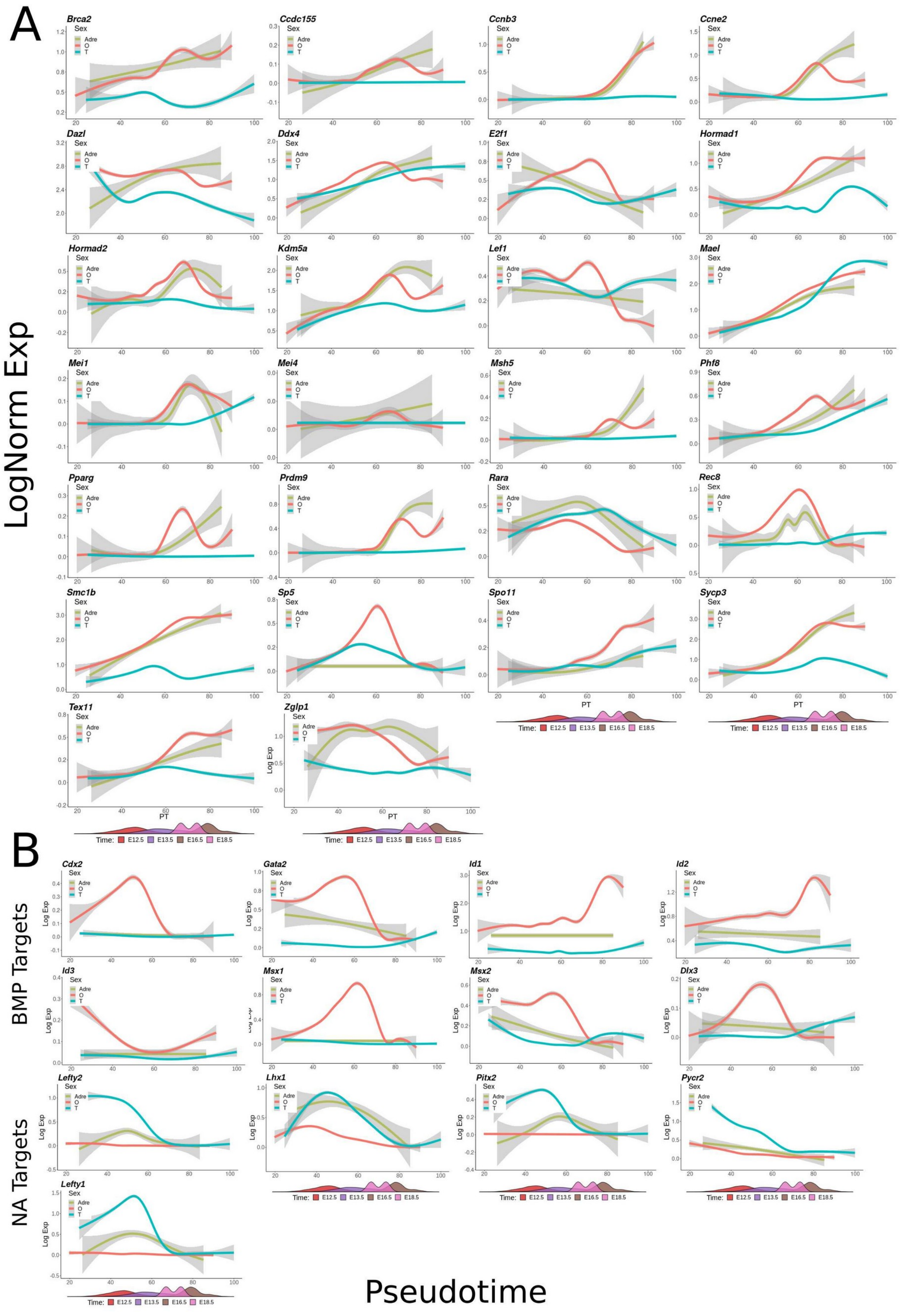
A
LogNorm Exp
Brca2
Ccdc155
Ccnb3
Ccne2
Dazl
Ddx4
E2f1
Hormad1
Hormad2
Kdm5a
Lef1
Mael
Mei1
Mei4
Msh5
Phf8
Pparg
Prdm9
Rara
Rec8
Smc1b
Sp5
Spo11
Sycp3
Tex11
Zglp1
Sex
Adre
O
T
PT
Time: E12.5 E13.5 E16.5 E18.5
B
BMP Targets
Cdx2
Gata2
Id1
Id2
Id3
Msx1
Msx2
Dlx3
NA Targets
Lefty2
Lhx1
Pitx2
Pycr2
Lefty1
Log Exp
Pseudotime

**Figure S4**

**Legend Figure S4** (related to Figure 4)**. Smoothed expression curves of genes involved in meiosis, oogonia and spermatogonia differentiation process in gonadal and ectopic germ cells.** The solid line represents the smoothed expression curves of the gene in the cells ordered by pseudotime, and the fade band is the 95% confidence interval of the model.

#### Supplementary experimental procedures

##### Data processing with the Cell Ranger package, cell selection and in-house quality controls

For cell-associated barcode selection, we computed the knee point and the inflection point of the ranked barcodes distribution plot. The local minimum between these points on the density curve (density base R function, bw=500, n=4096) of the UMI number per barcode was detected using the quantmod package. This local minimum was used as a threshold for cell-associated barcode selections. When no local minimum could be detected between the two points, the nearest local minimum was used. Quality controls regarding abnormal mitochondrial or ribosomal content, UMI number, detected gene numbers, unmapped reads and putative doublet identification (Scrublet) were performed, but no data filtering was applied as no important anomalies were detected.

##### Generation of a single-cell transcriptional atlas of germ cells sex determination and differentiation

The transcriptomes of the individual cells were sequenced at the depth of ~150,000 reads/cell. After barcode filtering based on the unique molecular identifiers (UMI) distribution, we obtained 92,267 cells. It included 14,904 cells from E10.5, 16,581 cells from E11.5, 19,551 cells from E12.5, 25,012 cells from E13.5, and 16,219 cells from E16.5. Among the 52,463 XY cells and the 39,804 XX cells, the median number of UMIs was 17,493 and 17,655 and the median number of detected genes was 4,802 and 4,658, respectively.

##### Clustering and UMAP visualization

To cluster cells we first selected all genes detected in more than 50 cells, and performed ICA (n_comps=100, Seurat) on log normalized values. To assess for batch effect, we built a nearest neighbor graph using BBKNN function (BBKNN package).

To classify the cell populations present in the developing testis and ovary, we selected all genes detected in more than 50 cells (21,103 genes), log normalized their expression and ran Independent

Component Analysis (ICA) on these (100 components). ICA extracts non Gaussian components from data, allowing a better discrimination of small cell populations in heterogeneous dataset. We set the number of ICs to 100 as we expected to have less than 100 different populations in our total dataset. Another advantage of ICA is its good performance with nearly no gene filtering, such as highly variable genes selection. A neighbor graph corrected for batch between replicates was computed using BBKNN (Park et al., 2018) with default values and n_neighbors_within_batch = 3 and the ICA as input (all components). Louvain clustering was then applied with resolution 1. We obtained 50 cell clusters combining different embryonic stages and sexes (**Fig. S1A**). As expected, the clusters of cells from early stages (E10.5 and E11.5) tended to include XX and XY cells while at later stages cell clusters were usually sex-specific, reflecting the emergence of more differentiated cell populations. As expected, biological duplicates isolated from independent pregnancies represented equivalent developmental stages and sex and clustered tightly together (**Table S3**). Cells from clusters 6, 11, 13, 25, 28, 29, 39, 41 and 48 were identified as germ cells and selected for all subsequent analysis.

**Cell lineage reconstruction identifies the dynamics of gene expression during XX and XY germ cell sex determination**

To order cells along a pseudotime, we used ordinal regression modeling (Telley et al., 2018; Teo et al., 2010) using prior knowledge about the developmental stage of each cell capture. Ordinal regression modeling is similar logistic regression classifiers but allows to take into account the order of the classification categories, here the embryonic time. It therefore allows the generation of a continuous score predicting the belonging of cell to one or another embryonic time.

We trained the model using a ten fold cross-validation schema on the whole expression matrix with a correction for unequal numbers of XX and XY cells. The trained model outputs a weight for each gene according to its ability to predict the embryonic time of each cell in an ordered manner. We then predicted a continuous score for all cells using the weights of the 100 top weighted genes to avoid over fitting of the model.

To determine which genes were important for sex specific progression of the cells along time, we trained two distinct models on XX and XY cells and selected the 2,000 top-weighted genes and merged the two lists of genes. We obtained 3,013 unique genes, of which 987 were common to both XX and XY lineages, 1,013 were specific to XX germ cells, and 1,013 were specific to XY germ cells (**Fig. S2A,B and Table S1**)

**Gene regulatory network generation**

GRN were generated using pyScenic package (Aibar et al., 2017). First adjacencies were generated using the grnboost function (Moerman et al., 2019) with the whole expression matrix of all the germ cells as input and the transcription factor list provided with the package.

Next, modules were generated, associating groups of genes to a putative regulator using modules_from_adjacencies function. Finally, the obtained modules were pruned on the basis of the presence or absence of the master regulator binding site footprint in the neighborhood of the TSS of each regulated gene. AUCell was then ran on the obtained regulons to infer its activity within each cell.

**Heatmaps and expression curves**

Heatmaps were generated using R (packages pheatmap  and heatmaps, smoothing was performed using smoothheatmap function on log-normalized expression levels for genes and AUC enrichment values for regulons. Expression curves were generated using ggplot2 geom_smooth function with method “gam” on log normalized expression levels.

**Velocity analysis**

To generate spliced and unspliced counts data, the velocyto.py script from velocyto package (La Manno et al., 2018) was called on each bam file with aforementioned reference genome annotation.

Spliced and unspliced counts were normalized by dividing all UMI counts of a cell by the total counts in the cell, multiplying the result by the median count number in all cells and applying log1p function. We then used scvelo package for all subsequent analysis. Moments for velocity estimation were generated using pp.moments function with previously generated independent components. Gene specific velocity and velocity graph were then computed following workflow provided on scvelo documentation with default arguments. UMAP plotting of a specific gene with spliced and unspliced counts were generated using Ms and Mu layers (kNN pooled expression) respectively. For smoothed expression curves, log normalized spliced counts were multiplied by the velocity_gamma parameter generated for each gene by the tl.velocity function.

**Ectopic adrenal germ cells analysis**

The analysis was performed using aforementioned steps: log normalization, ICA, neighbor graph, and clustering with the same parameters. Germ cell clusters were selected with the same 10 markers genes. For pseudotime ordering of the cells, a model was trained on gonadal cells only and the pseudotime for adrenal germ cells was predicted from it.
